# Curiosity satisfaction increases event-related potentials sensitive to reward

**DOI:** 10.1101/2023.05.17.540372

**Authors:** Tim Rueterbories, Axel Mecklinger, Kathrin C. J. Eschmann, Jordan Crivelli-Decker, Charan Ranganath, Matthias J. Gruber

## Abstract

Successful learning depends on various factors such as depth of processing, motivation, or curiosity about information. A strong drive to learn something or the expectation of receiving a reward can be crucial to enhance learning. However, the influence of curiosity on the processing of new information and its similarity with reward processing is not well understood. This study examined whether states of curiosity influence specific event-related-potentials (ERPs) associated with reward processing and whether these ERPs are related with later memory benefits. In an initial screening phase, participants indicated their curiosity and confidence in prior knowledge about answers to various trivia questions. In a subsequent study phase, we targeted different time windows related to reward processing during the presentation of trivia answers containing the Reward-Positivity (RewP; 250 - 350 ms), the P3 (250 - 500 ms), and the Late-Positive-Potential (LPP; 600 - 1000 ms). In a following surprise memory test, we found that participants recalled more high- than low-curiosity answers. The RewP, P3 and LPP showed greater positive mean amplitudes for high compared to low curiosity, reflecting increased reward processing. In addition, we found that the RewP and the P3 showed more positive mean amplitudes for later recalled compared to later forgotten answers, but curiosity did not modulate this encoding-related results. These findings support the view that the satisfaction of curiosity resembles reward processing, indicated by event-related-potentials.

## Introduction

Why did you start reading this paper? Maybe because you stumbled across it by accident, maybe because you have to, or because you are curious about its content. Curiosity is an intrinsic motivation that drives us to collect new information and has been shown to enhance memory formation (Gruber & Ranganath, 2019; Galli et al., 2018; Kidd & Hayden, 2015; Loewenstein, 1994). For example, if someone asks you “What is the longest river in the European Union?”, you might be curious and try to come up with the correct answer. In the last decade, initial research on states of curiosity have targeted the neural underpinnings of the *elicitation* of curiosity (e.g., when you read a question without knowing the answer; Poh et al., 2022; Lau et al., 2020; Oosterwijk et al., 2020; Ligneul et al., 2018; Gruber et al., 2014; Kang et al., 2009), but the literature still lacks findings as to what specific processes take place during the *satisfaction* of curiosity (e.g., when you receive the answer “Danube”).

Most concepts about states of curiosity have in common that the awareness of a knowledge gap leads to increased exploration for new information (Metcalfe et al., 2020; Gruber & Ranganath, 2019; Loewenstein, 1994; Berlyne, 1954). New information satisfies curiosity, reduces uncertainty, can improve predictions in the future, and can be described as an unconditioned reward stimulus (Kang et al., 2009). Therefore, curiosity reflects an intrinsically anticipated expectation of reward that intensifies the processing of upcoming information (Gruber & Ranganath, 2019; Marvin & Shohamy, 2016; Kidd & Hayden, 2015; Lowenstein, 1994). Investigating event-related-potentials (ERPs) during the satisfaction of curiosity may possibly explain how curiosity influences the processing of information, what similarities exist regarding to reward processing, and how curiosity satisfaction modulates memory formation.

There is evidence showing that the neural correlates of curiosity resemble those of reward anticipation (Lau et al. 2020; Oosterwijk et al. 2020; Gruber et al., 2014) and reward processing (Ligneul et al., 2018; Jepma et al., 2012). For example, examining curiosity satisfaction with ambiguous visual images, Jepma et al. (2012) showed that the resolution of the images was associated with increased activity of the striatum, which has been related to reward processing. Furthermore, investigating effects of intrinsic (curious) and extrinsic (rewarding) motivation on memory, Duan et al. (2020) found that curiosity-driven memory formation was associated with the ventral striatal reward network and the fronto-parietal attention system, whereas extrinsic-driven memory effects were correlated with deactivation in parietal midline regions. Therefore, it is not clear whether reward and curiosity are really the same processes. To investigate this question, we therefore assumed that curiosity satisfaction should be similar with reward processing and predicted that reward-related ERPs should also be modulated by states of curiosity.

As the answer to a question can be seen as an unconditioned reward stimulus, we chose to investigate the Reward Positivity (RewP; 250 ms – 350 ms; Proudfit, 2015), also known as Feedback-Related-Negativity (FRN; Höltje & Mecklinger, 2020; Peterburs et al., 2016; Cohen & Ranganath, 2007). According to Proudfit (2015, p. 449), the FRN “reflects a reward-related positivity that is absent or suppressed following non-reward”. Since in this study we assume that the answer represents a reward, we favour the interpretation as RewP. This component is assumed to reflect a better outcome than expected, an increase in dopaminergic signals, and reward processing at fronto-central sites (Glazer et al., 2018; Heydari & Holroyd, 2016; Proudfit et al., 2015; Sambrook & Goslin, 2015). In addition, the RewP seems to be correlated with neural activity in reward-mediating regions such as the ventral striatum, anterior cingulate cortex, and medial prefrontal cortex (Becker et al., 2014). We hypothesized that when curiosity is high, a new information should be perceived as more relevant, reflecting a greater reward, and should therefore result in a larger mean amplitude of the RewP.

After early reward processing reflected by the RewP, the P3 (P300/FB-P3; further: P3; 250 ms – 500 ms; Polich, 2007) could be also manipulated due to high- and low-curiosity levels. The P3 is usually found at central and parietal electrodes and assumed to reflect the allocation of neural resources based on reward effects, stimulus relevance, as well as context updating in working memory (Van Petten & Luka, 2012; San Martin, 2012). In addition, it displays attention-driven categorization and increased processing of new information due to motivational salience. The P3 amplitude is larger if new information is infrequent or a reward is greater than expected (Hajcak et al., 2006). For high curiosity, it is assumed that there should be a greater mean amplitude of the P3 during curiosity satisfaction because of higher motivational relevance for the updating of the stimulus context, increased attention, and a subjective greater reward than for low curiosity (Donaldson et al., 2016; Polich, 2007).

The Late-Positive-Potential (LPP; 600 ms – 1000 ms; Glazer et al., 2018) is a centro-parietal positive-going ERP component that has been associated with the processing of emotional stimuli as well as extended cognitive and attentional processing based on reward expectancy and magnitude (Hajcak & Foti, 2020; Glazer et al., 2018; Meadows et al., 2016; Gable et al., 2015; Weinberg et al., 2013; Schupp et al., 2000). Furthermore, the LPP appears to be manipulated by the significance of a stimulus, which is determined by the activation of motivational appetitive systems (Bradley, 2009). Based on these findings, we expect the mean amplitude of the LPP to be more positive for high-than for low-curiosity answers.

To generate states of high and low curiosity, we used the so-called trivia paradigm and adopted an identical experimental design as in a previous functional magnetic resonance imaging (fMRI) study from our lab (Gruber et al., 2014). In an initial screening phase, participants were presented with general knowledge questions, and were asked to indicate their curiosity and confidence in prior knowledge about the answer. In the following study phase, participants were presented with a selected set of trivia questions from the screening phase and a few seconds later with the associated answers (i.e., the satisfaction of curiosity). ERPs were computed time-locked to the onset of the answers. After a five-minute break, participants were asked to recall the answers in a surprise memory test.

Based on previous findings (e.g., Poh et al., 2022; Wade & Kidd, 2019; Marvin & Shohamy, 2016; Gruber et al, 2014; Kang et al., 2009), we predicted memory performance to be better for answers associated with high compared to low curiosity. Furthermore, we expected that ERPs recorded during the presentation of the answer should differentiate between remembered and forgotten ones. To address this question, we used the memory test results to investigate subsequent memory effects (SMEs), for which the ERPs are sorted according to whether the answers are later remembered or forgotten. In general, subsequently remembered items should show a more positive ERP amplitude during encoding than those that have been forgotten (for review see Mecklinger & Kamp, 2023; Cohen et al., 2015). Accordingly, we predicted more positive amplitudes for remembered answers. Furthermore, we also explored the interaction between curiosity and memory in relation to the ERP amplitudes.

## Materials & Methods

### Participants

Thirty healthy young adults participated in the experiment A required sample size of N = 24 was determined with a power analysis for a repeated-measures ANOVA (G*Power, Version 3.1.9.7.; Faul et al., 2009), i.e., the ERP amplitude difference between high and low curiosity, based on the assumption of a medium effect size according to Cohen’
ss “f” (1988), f = 0.25, α = 0.05, 1-β = 0.8. Eight participants had to be excluded due to either poor memory performance (in at least one condition fewer than eight memorized answers) or incomplete EEG or memory data. Since our final data set was slightly below the targeted sample size, we additionally calculated the Bayes Factor for all non-significant results. This tests, whether potential null results are due to inconclusive data or whether there is actual support for a null hypothesis (Dienes, 2014). The final sample included N = 22 participants (11 female, 11 male, age range = 18-30 years, *M* = 21.23 years). One participant was left-handed and twenty-one right-handed. All participants had normal or corrected-to-normal vision and were English native speakers. All participants were students at University California (UC) Davis and received money or course credits for their attendance. The UC Davis Institutional Review ethics committee approved the experiment.

### Material

The questions and their matching answers for the screening and study phase were randomly drawn from a pool of 381 items (https://osf.io/he6t9/), which in turn were acquired from online resources about general knowledge questions. In addition, care was taken to select from as many different subject areas as possible (i.e., sports, food, science, nature, TV/movies, music, history etc.). The stimuli were always presented centred in black on a grey background.

### Procedure

The experiment was programmed with the Cogent 2000 Toolbox (Wellcome Laboratory of Neurobiology) and carried out in the Center for Neuroscience at UC Davis. The experiment is a version of the trivia paradigm (cf., Kang et al., 2009) and was divided into a screening, study, and test phase (for an almost identical task procedure see: Gruber et al., 2014; for an overview of the procedure, see Figure 1).

**Figure 1.**
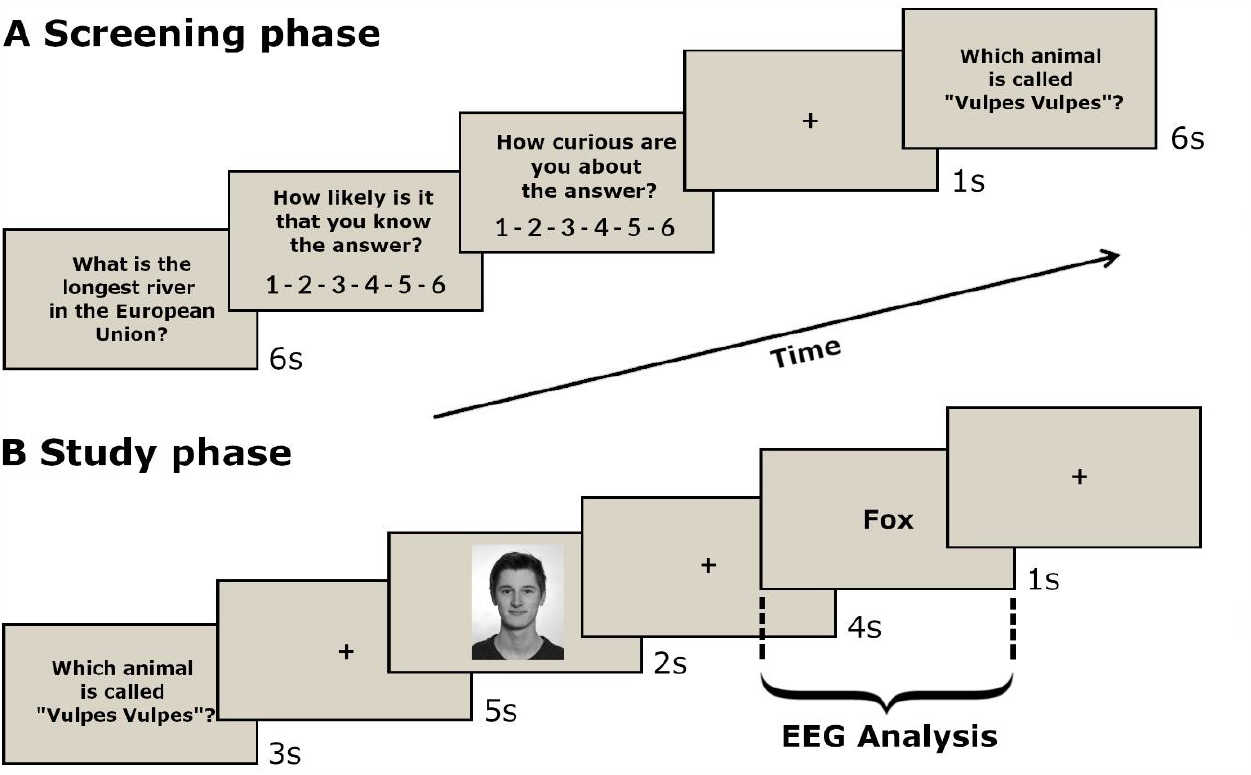
Experimental Procedure. **(A)** In the screening phase, trivia questions were presented and evaluated by participants according to their knowledge confidence and perceived level of curiosity without reading the corresponding answers. **(B)** In the following study phase, participants are presented with a subset of these questions again and eleven seconds later with their corresponding answers. Between questions and answers, emotionally neutral faces were presented. The instruction of the participants was to “always try to anticipate the possible answer and give a response (“YES” or “NO”) whether the person can help you”. The EEG was recorded during the study phase and ERPs were computed time-looked to the onset of the answer. **(C)** At last, participants were asked to recall all answers in a surprise memory test (not shown).

In the screening phase, randomly chosen questions were presented for six seconds and evaluated by participants according to their knowledge confidence and perceived level of curiosity. This intra-individual rating of the questions was necessary because curiosity levels for each trivia question and its respective answer differ between individuals. First, participants were asked to indicate how confident they were that they knew the answer to the trivia question (1-6; 1 = ‘‘I am confident that I do not know the answer’’ and 6 = ‘‘I am confident that I know the answer’’). As a second judgment, participants indicated their level of curiosity about the answer (1-6; 1 = ‘‘I am not interested at all in the answer’’ and 6 = ‘‘I am very much interested in the answer’’). Questions were presented until 56 items could be attributed to high (curiosity rating 4-6) and 56 to low curiosity (curiosity rating 1-3). However, if the participants had given a “6” for the confidence rating, the question was not included, because it was assumed that participants already knew the answer. While participants were completing the screening phase, experimenters were attaching the EEG cap to the participant and were preparing all electrodes in order to record the EEG. The screening phase was followed by a short rest period (approx. 5 min) during which participants looked at a fixation cross.

In the study phase, 112 previously selected questions were presented again (3s) and after an anticipation period (11s; the time between question presentation offset until answer presentation onset), the matching answers were displayed (1s). Six trials in each condition (∼ 10%) were catch trials, during which a sequence of letters “xxxxx” were shown as answer, in order to keep participants’ attention high. To create consistency to our prior work using the identical paradigm in a fMRI study (see Gruber et al., 2014) emotionally neutral faces (2s) were presented during the anticipation period. The instruction of the participants was to “always try to anticipate the possible answer and give a response (“YES” or “NO”) whether the person can help you”. Since our manuscript focuses on the satisfaction of curiosity (i.e., during answer presentation) and thus on specific ERPs that previously have been shown to be sensitive to reward processing, the analysis of the faces during the anticipation of curiosity were not part of the present study.

After a further five-minute rest period, during which participants looked at a fixation cross, a surprise memory test was carried out in the test phase. Participants were given a randomized list with all trivia questions from the study phase. They were encouraged to type in as many answers as possible within twenty minutes without guessing the answers. Participants also took part in a recognition memory test for all previously encoded face images, which are not part of this investigation (see Gruber et al., 2014 for face analysis). Each participant took part in each phase once and received money or course credits after the test phase.

### EEG recording and processing

During the study phase, the EEG was recorded with the ActiveTwo EEG recording system (Biosemi, Neurospec) at 1024 Hz from 64 Ag/AgCl scalp electrodes, which were arranged according to an extended version of the international 10-20 electrode system (Jasper, 1958). The electrodes were offline re-referenced to the averaged mastoid electrodes. The vertical and horizontal electrooculogram was recorded from four electrodes placed above and below the left eye and at the canthi of the left and right eye.

The analysis of the EEG data was performed using MATLAB (MathWorks, Inc.) and the toolboxes EEGLAB (Delorme & Makeig, 2004) and ERPLAB (Lopez-Calderon & Luck, 2014). Electrodes were referenced to the average of the left and right mastoid electrodes. EEG data were down sampled to 500 Hz and bandpass filtered at 0.1-40 Hz using a second order Butterworth filter. The epochs started at -200 ms before stimulus presentation and ended at 1000 ms thereafter. In addition, data segments outside of the area of interest were discarded and an independent component analysis (ICA) was applied to correct for artifacts. Segments, which were still associated with ocular and noise artifacts, were rejected manually for each participant. Additionally, segments containing artifacts were removed based on the following criteria: The maximum voltage threshold was set between -150 μV and +150 μV, the maximal permissible difference of values at 150 μV during intervals of 200 ms, and the maximum allowed voltage difference between two time points was 30 μV. To exclude data with no signal, trials which did not exceed the limits -0.7 μV and +0.7 μV within a period of 200 ms were discarded. On average, 83.93% of trials were retained (*M* = 94, trial range = 40-110, high curiosity = 46.89, low curiosity = 47.11) and 16.07% rejected (*M* = 18, range = 2-72, high curiosity = 9.11, low curiosity = 8.89). These cleaned and pre-filtered data were pruned and merged based on curiosity level and memory performance. We calculated the mean amplitudes for the components of interest over all trials for each participant.

The RewP is typically measured at fronto-central sites and thus activity at electrodes FC1, FCz, and FC2 in the time range of 250 ms – 350 ms was investigated (Proudfit et al., 2015). The P3 is usually found at central and parietal electrodes, which is why activity at electrodes Cz, CPz, and Pz between 250 - 500 ms was analysed (Polich, 2007). The LPP is most pronounced at centro-parietal electrodes and hence activity at electrodes CP1, CPz, and CP2 in the time range of 600 - 1000 ms after stimulus onset was explored (Glazer et al., 2018; Gable et al., 2015). The time window of the LPP can also be broader, but our time window was limited to 1000 ms.

### Statistical analyses

The statistical analyses were performed using R Studio software (R Core Team, 2019). The significance level was set to α = .05 and *t*-tests were one-tailed based on specific hypotheses. For the behavioral analyses, we used dependent *t*-tests and regression models with curiosity and confidence as predictors of memory. The ERP data was analysed using 2 (curiosity: high, low) × 2 (memory: remembered, forgotten) repeated-measures ANOVAs, with the values averaged across the electrodes. To address the question of sufficient evidence for non-significant results, we additionally computed the Bayes Factor (Dienes, 2014). For interpretation, we used the widespread labels by Jeffreys (1961): BF_10_ < 0.3 as substantial evidence for H_0_ and BF_10_ > 3 as sub. evidence for H_1_. As measures of effect sizes, partial eta squared 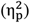 is reported for ANOVA and Cohen’s *d* (Rasch et al., 2021; Cohen, 1988) was calculated for dependent *t*-tests.

## Results

### Behavioural results

The analysis of memory performance showed a significant mean recall difference between high- (*M* = 53.32%, *SD* = 15.57%) and low-curiosity answers (*M* = 37.15%, *SD* = 16.19%), *t* (21) = 7.37, *p* < .001, *d* = 1.57, which replicates the previously found high-curiosity related memory benefit (Kang et al., 2009; Gruber et al, 2014). The overall memory performance was *M* = 42.00% (lowest = 16.96%; highest = 67.86%) and the curiosity benefit was *M* = 16.17% (high-curiosity answers – low-curiosity answers). In addition, a follow-up behavioural analysis using the 6-scale curiosity ratings to predict memory was conducted to check whether a linear relationship exists between both variables and thus the classification to high and low curiosity can be supported. The logistic regression shows that as curiosity increases, memory performance also increases continuously (*β* = 0.21, *SE* = 0.03, *z* = 8.26, *p* < .001). Accordingly, analysing curiosity with a two-level factor (high and low) is an adequate approach for the ERP analyses (see Figure 2).

**Figure 2.**
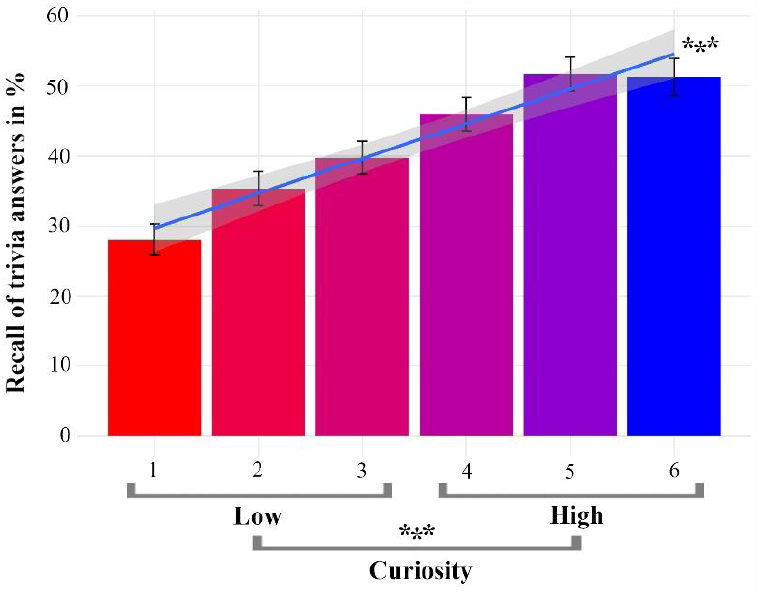
Curiosity-related memory benefit. Better memory performance for trivia answers in the high-compared to the low-curiosity condition. Error bars indicate the standard error of the mean. The line represents a linear model between curiosity and memory. * p < .05, ** p < .01, *** p < .001

Moreover, calculating the confidence ratings for high (*M* = 2.26, *SD* = .45) and low curiosity (*M* = 1.50, *SD* = .34) revealed a significant difference, *t* (21) = 7.64, *p* < .001, *d* = 1.63, indicating that participants were more confident they knew the answers in relation to high curiosity. A correlation between curiosity and confidence was significant, with r = .35, p <.001, showing that 12.25% of variance caused by curiosity can be explained by confidence. These analyses show that curiosity and confidence in prior knowledge are partly related, but most of their variance exists independently.

Most importantly, to make sure that our curiosity effects on learning remain regardless of confidence in prior knowledge, we ran a follow-up logistic regression analysis. Controlling for each other, both z-scored curiosity (β = 0.26, SE = 0.05, z = 5.69, p < .001) and z-scored confidence (β = 0.26, SE= 0.05, z = 5.45, p < .001) were significant predictors of memory. Critically, there was no interaction between curiosity and confidence in predicting memory (β = -0.08, SE = 0.05, z = -1.66, p = .096; *BF*_*10*_ = *0*.*05*). Our behavioral results, along with previous findings in the literature (Wade & Kidd, 2019; Stare et al., 2018), suggests that although confidence has an impact on memory, the effect of curiosity remains independent of confidence in prior knowledge.

### Electrophysiological results Reward Positivity (250 – 350 ms)

To investigate reward-related processes during the presentation of trivia answers, the RewP was analysed in a 2 (curiosity: high, low) × 2 (memory: remembered, forgotten) repeated-measures ANOVA. This analysis yielded a significant main effect for curiosity, *F*(1, 21) = 6.29 *p* = .020, η_p_ ^2^ = .23, and for memory, *F*(1, 21) = 6.43, *p* = .019, η_p_ ^2^ = .23 (Figure 3). Notably, a 2-way interaction between curiosity and memory was not significant, *F*(1, 21) = 0.24, *p* = .63, η_p_ ^2^ = 0.01; *BF*_*10*_ = *0*.*15*. The RewP amplitude was more positive for high-curiosity (*M* = 3.26, *SD* = 3.54) compared to low-curiosity answers (*M* = 2.07, *SD* = 3.56; *t* (21) = 2.51, *p* = .010, *d* = 0.54) as well as for remembered answers (*M* = 3.41 *SD* = 3.51) compared to forgotten answers (*M* = 1.91, *SD* = 3.78; *t*(21) = 2.54, *p* = .009, *d* = 0.54). It can be assumed that the greater RewP amplitude for high curiosity answers indicates increased reward-related processing of the answers in comparison to low curiosity. Furthermore, a greater amplitude of the RewP was predictive for better memory, but we did not find that increased reward processing benefitted later memory, that is, there was no interaction between curiosity and memory for the RewP.

**Figure 3.**
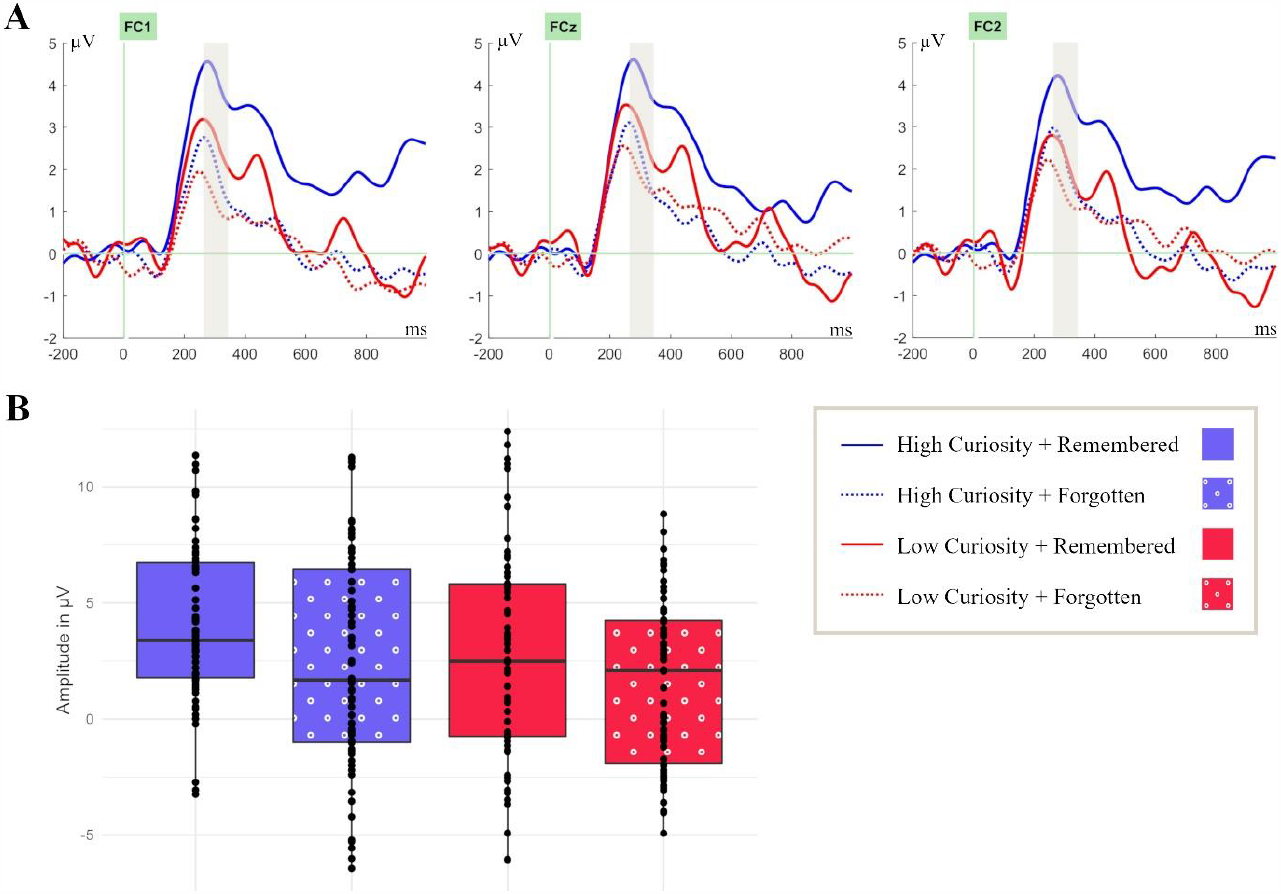
Curiosity and memory independently modulate the Reward Positivity (RewP) component during answer presentation at fronto-central electrode sites. **(A)** ERPs were time-locked to the onset of the answers in the study phase at electrode sites FC1, FCz, and FC2 in the coloured time window of 250 – 350 ms. **(B)** Box plots are bounded by the first and third quartiles and the black line represents the median. The points correspond to the measures for individual participants.

### P3 (250 – 500 ms)

For examining the allocation of neural resources based on reward effects, stimulus relevance, and context updating in working memory, the P3 was analysed in a 2 (curiosity: high, low) × 2 (memory: remembered, forgotten) repeated-measures ANOVA. Similarly, to the findings of the RewP, this analysis yielded a significant main effect for curiosity, *F* (1, 21) = 4.70, *p* = .042, η_p_ ^2^ = .18, as well as for memory, *F* (1, 21) = 6.70, *p* = .017, η_p_ ^2^ = .24. (Figure 4). The 2-way interaction between curiosity and memory was not significant, *F* (1, 21) = 0.26, *p* = .61, η_p_ ^2^= 0.01; *BF*_*10*_ = 0.16. The amplitude was more positive for high-curiosity (*M* = 2.83, *SD* = 3.99) compared to low-curiosity answers (*M* = 1.76, *SD* = 3.31; *t* (21) = 2.17, *p* = .021, *d* = 0.46) as well as for remembered answers (*M* = 2.92 *SD* = 3.66) compared to forgotten answers (*M* = 1.67, *SD* = 3.65; *t* (21) = 2.59, *p* = .008, *d* = 0.55). Most relevantly, the greater P3 amplitude for high curiosity suggests increased stimulus relevance as well as more pronounced context updating in working memory. In addition, as with the RewP, a greater amplitude of the P3 was predictive for better memory.

**Figure 4.**
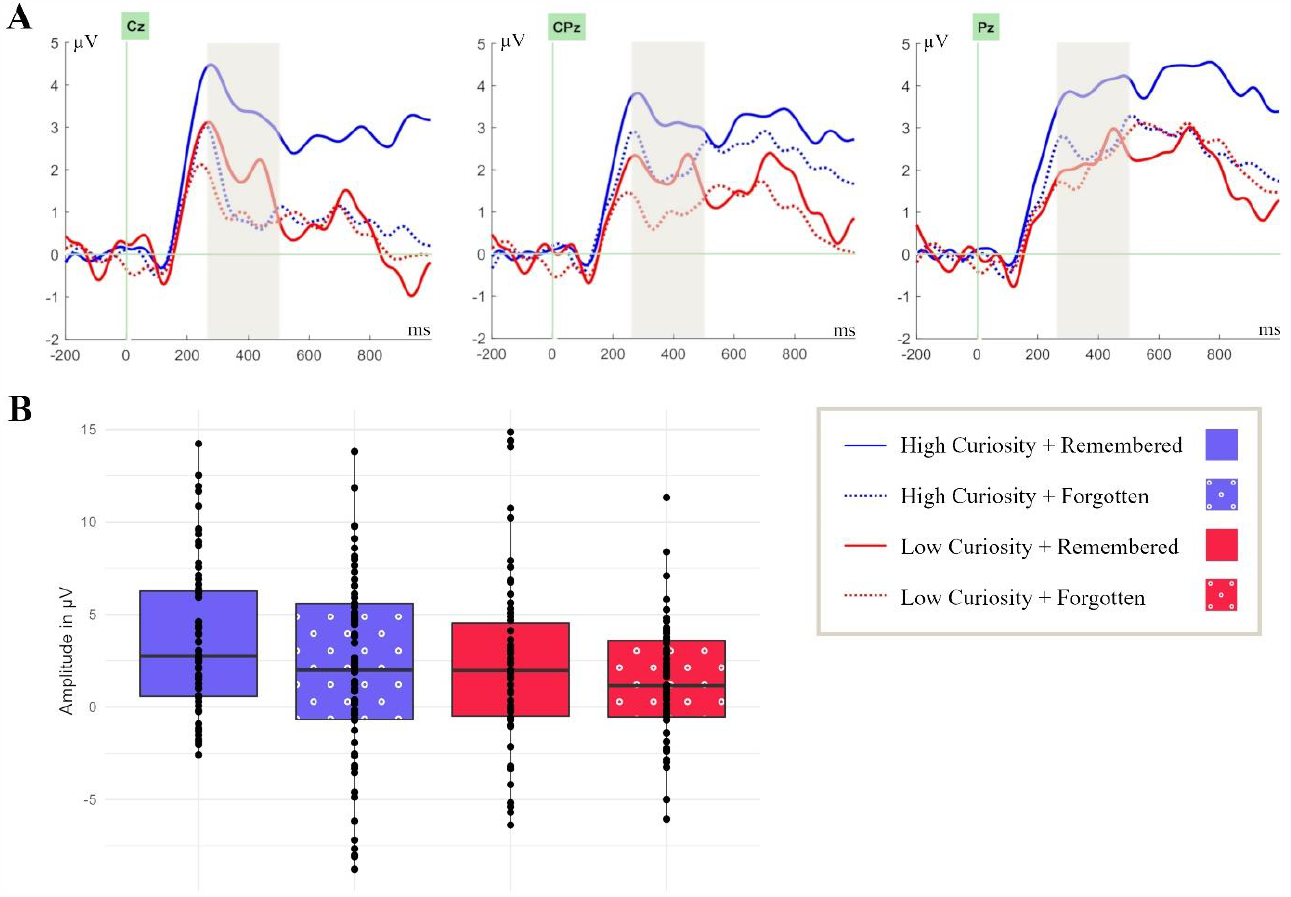
Curiosity and memory independently modulate the P3 component during answer presentation at central and parietal electrode sites. **(A)** ERPs were time-locked to the onset of the answers in the study phase at electrode sites Cz, CPz, and Pz in the coloured time window of 250 - 500 ms. **(B)** Box plots are bounded by the first and third quartiles and the black line represents the median. The points correspond to the measures for individual participants.

### Late-positive-potential (600 – 1000 ms)

The analysis of extended cognitive and attentional processing based on reward expectancy and magnitude was implemented in a 2 (curiosity: high, low) × 2 (memory: remembered, forgotten) repeated-measures ANOVA on the LPP. This analysis yielded a significant main effect for curiosity, *F* (1, 21) = 5.16, *p* = .034, η_p_ ^2^ = .20, but not for memory, *F* (1, 21) = 0.83, *p* = .37, η_p_ ^2^ = .04, *BF*_*10*_ = 0.27 (Figure 5). There was no 2-way interaction between curiosity and memory, *F* (1, 21) = 1.26, *p* = .27, η_p_ ^2^ = 0.06; *BF*_*10*_ = 0.26. The amplitude was more positive for high-curiosity (*M* = 2.66, *SD* = 5. 40) compared to low-curiosity answers (*M* = 1.32, *SD* = 4.61; *t* (21) = 2.27, *p* =.017, *d* = 0.48). The greater LPP amplitude for high-compared to low-curiosity answers is assumed to indicate extended cognitive and attentional processing based on reward magnitude.

**Figure 5.**
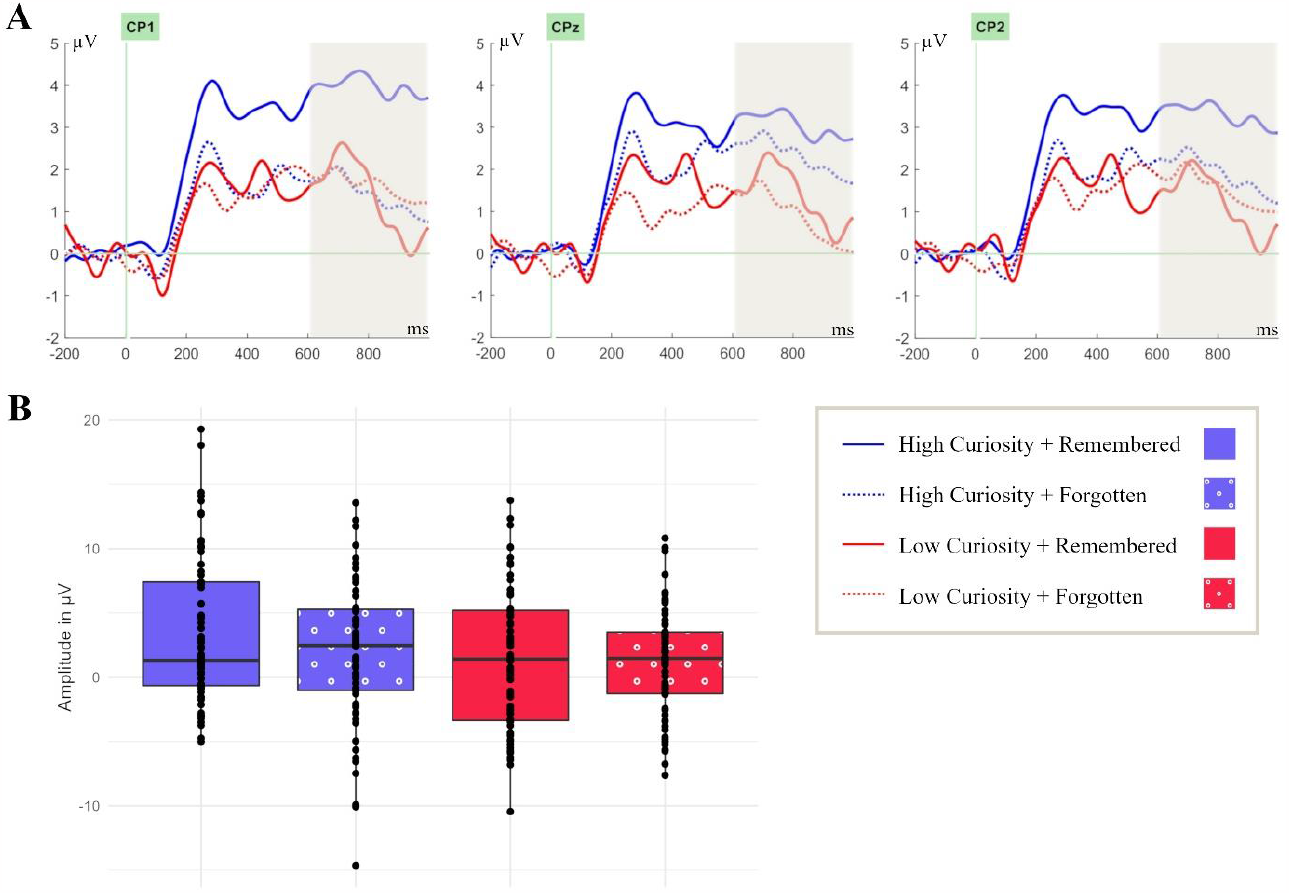
Only curiosity (but not memory) modulates the Late-Positive-Potential (LPP) during answer presentation at centro-parietal electrode sites. **(A)** ERPs were time-locked to the onset of the answers in the study phase at electrode sites CP1, CPz, and CP2 in the coloured time window of 600 - 1000 ms. **(B)** Box plots are bounded by the first and third quartiles and the black line represents the median. The points correspond to the measures of individual participants.

## Discussion

The present paper addressed the question of whether curiosity satisfaction modulates specific ERPs that are sensitive to reward. In the initial screening phase, participants were presented with general knowledge questions and were asked to indicate their level of curiosity and confidence of one’s own prior knowledge about the respective answer. In a subsequent study phase the influence of curiosity levels on reward-related ERPs was examined via the RewP, P3, and LPP. At the end of the experiment there was an unannounced memory test in which the subjects had to repeat the previously presented answers to the trivia questions.

Behaviourally, we found better memory performance for high-curiosity in contrast to low-curiosity answers. In the same study setup, Gruber et al. (2014) found similarly large differences between high- and low-curiosity memory performance. In general, the reported memory benefit for high curiosity is in line with the findings of several other studies (Poh et al., 2022; Fandakova & Gruber, 2021; Lau et al., 2020; Galli et al., 2018; Stare et al., 2018; Marvin & Shohamy, 2016; Gruber et al, 2014; Kang et al., 2009). Our analyses also showed that curiosity and confidence in one’s own prior knowledge influence memory independently and tend to share only a small proportion of variance. It can therefore be assumed that the results of the ERP analyses can be attributed to the different levels of curiosity independent of confidence.

ERP analysis aimed at assessing differences in the online processing of high-compared to low-curiosity answers and processes predictive of successful memory formation. We found that the amplitudes of the RewP, P3, and LPP were more positive in the high-than in the low-curiosity condition. Furthermore, the RewP and P3 (but not the LPP) showed greater positive amplitudes for later remembered compared to later forgotten answers. No interaction between curiosity levels and subsequent memory were found for the examined ERP components. However, this interaction results should be viewed with caution due to the exploratory approach. In general, the results suggest that the satisfaction of curiosity via new information resembles reward processing.

### Reward Processing

Firstly, we assumed that information – here the answer to a trivia question – represents a rewarding stimulus that reduces uncertainty and closes a knowledge gap (Loewenstein, 1994). Several behavioural and neuroimaging studies support the view that states of high curiosity and extrinsically motivated reward depend on similar mechanisms and activate overlapping brain areas (FitzGibbon et al., 2020; Marvin & Shohamy, 2016; Gruber et al., 2014; Kang et al., 2009; Bromberg-Martin & Hikosaka, 2009). Moreover, the examined RewP is seen as an indicator of reward processing (Anderson et al., 2017; Heydari & Holroyd, 2016; Sambrook & Goslin, 2015). Accordingly, the discovered differences of the RewP amplitude based on curiosity levels can be seen as a neurophysiological correlate of reward processing, where answers with high curiosity triggered greater reward responses.

In addition, the Prediction, Appraisal, Curiosity, and Exploration (PACE) framework (Gruber & Ranganath, 2019) suggests that higher expectation of reward and curiosity are accompanied by increased dopaminergic activity, which leads to facilitated hippocampus-based memory formation (Gruber & Ranganath, 2019; Smith et al., 2005; Wittmann et al., 2005; Schultz, 2002; Holroyd & Coles, 2002). Even though the present work does not allow to draw any direct conclusions about dopaminergic circuit activity, the identical experimental paradigm as in our previous fMRI study on curiosity (Gruber et al., 2014) permits to assume an increase in dopaminergic activity. Furthermore, the RewP can be an indication of dopaminergic signals itself (Baker & Holroyd, 2011). The origin of the dopaminergic activity echoed by the RewP has been localized in the basal ganglia and other reward-related brain areas and positive reward seems to be associated with increased medial prefrontal cortex (mPFC) activity (Becker et al., 2014). Moreover, van Lieshout et al. (2018) found increased mPFC activity during curiosity satisfaction and Murphy et al. (2021) investigated functional connectivity between the hippocampus and mPFC during answer presentation, which predicted the curiosity-related memory benefit. In summary, the increased positivity of the RewP in the high curiosity condition indicates, that the answers were perceived as more rewarding, which in turn seems to have strengthened reward-related processing.

Secondly, larger P3 and LPP amplitudes in the high curiosity condition may reflect increased allocation of neural resources and attention based on reward processing. A more positive amplitude of the P3 for high-compared to low-curiosity answers suggests that there was an enhanced allocation of neural resources and thereby increased attention (San Martin, 2012; Debener et al., 2005; Friedman et al., 2001). Suitable for this, the increased activity of the dopaminergic system (indicated by the RewP) can lead to more attention to upcoming rewards (Anderson et al., 2016; Hickey et al., 2010). First evidence for this was provided by fMRI studies that showed increased activity at high curiosity in frontal and parietal brain areas associated with attentional and cognitive control (Lieshout et al., 2018; Jepma et al., 2012). The modulation of the LPP also suggests that the perceived relevance of the information (reward magnitude) might have been greater during states of high curiosity due to the activation of appetitive motivational systems through curiosity (Hajcak & Foti, 2020). Synoptical, the greater positivity in the high curiosity condition of the P3 (Van Petten & Luka, 2012; San Martin, 2012) and the LPP (Weinberg et al., 2013; Schupp et al., 2000) can also be regarded as a marker of an increased reward processing.

Thirdly, the generally more positive amplitudes with high curiosity are in line with other studies examining answer processing after the presentation of trivia questions. Investigating the Tip-of-the-Tongue (TOT) phenomenon (see Metcalfe et al., 2017, Schwartz, 2006) – a subjective feeling associated with the state of high curiosity - Bloom et al. (2018) found an enhanced positivity during answer presentation at centro-parietal electrode sites, when participants reported to be in a TOT state. This study supports our conclusions that identifying a gap in knowledge can enhance processing of new information, as seen in the increased positive amplitudes of the ERPs.

In order to ensure consistency to our prior work using the identical paradigm in an fMRI study (see Gruber et al., 2014), we also adopted the time intervals of stimulus presentation. However, from the literature on feedback processing it is known that a long delay in presentation of a response (11 seconds in this study), can lead to a reduction of the RewP/FRN (Höltje & Mecklinger, 2018; Yin et al., 2018; Peterburs et al., 2016). For example, Yin et al. (2018) showed that the RewP decreases when feedback is delayed. At the same time, Höltje & Mecklinger (2018) were able to show that the FRN amplitude is reduced, but still reliable even with long delays (6,5s). Given that we found significant differences between curiosity conditions for RewP despite this longer response delay, this is evidence that curiosity might affect the RewP in a robust way.

Further, the instruction of the participants was to “always try to anticipate the possible answer”. Therefore, one cannot completely exclude the influence of expectation in this reward processing study. For example, the increased amplitude of the LPP (comparable with the P600 between 600 ms – 1000 ms; Kuperberg et al., 2020; DeLong et al., 2014) by high curiosity can potentially also be interpreted as increased integration effort of the answers based on expectancy. Late positive potentials have proven to be relevant EEG markers for the processing and integration of new information into context (Aurnhammer et al., 2021; Brouwer et al., 2017; Kos et al., 2012; Kuperberg, 2007). Since neither the contextual prior knowledge nor expectancy errors were collected in this study, this idea could be addressed in future research. Moreover, this also applies vice versa to studies that examine expectations, where feedback can also be rewarding. According to Wu & Zhou (2009), the FRN/RewP is more affected by expectancy related processing than the P300. As we found the curiosity effect on both components (RewP and P300), we take this as evidence that these effects reflect the influence of curiosity rather than feedback expectation.

### Memory effects

A SME was found for the amplitudes of the RewP and P3 without an interaction with curiosity. On the one hand, the difference in the amplitudes between remembered and forgotten words implies that the processes reflected by the RewP/P3 contribute to successful memory formation. On the other hand, the lack of an interaction does not allow for a direct connection between the curiosity modulation of the RewP/P3 and the behavioural curiosity-related memory benefits.

The SMEs could have been influenced by previous occurring processes, such as increased hippocampal activity or dopaminergic activity. It is important to note that Gruber et al. (2014) also did not find any significant interaction between curiosity-related brain activity and memory during answer presentation in the dopaminergic regions and the hippocampus. Critically, activation of dopaminergic areas and the hippocampus during curiosity elicitation but not satisfaction seems to support the curiosity-related memory benefit. During curiosity satisfaction, it seems that integration processes between hippocampus and large scale cortical networks (e.g., with the default mode network, Murphy et al., 2021) might contribute to the curiosity-related memory benefit. It is also conceivable that high curiosity levels might have enhanced the distinctiveness of events in a given processing context and the more distinctive items were more efficiently encoded and gave rise to an SME in the P3 time window (Otten & Donchin, 2000).

The absence of the SME for the LPP can have several reasons. On the one hand, it is possible that SMEs were too small to modulate the investigated ERP components significantly due to the sample size. On the other hand, SME effects for the LPP with onset latencies beyond 600 ms have typically been reported when intentional and elaborative encoding strategies emphasizing associative processing of multiple study features are required (Kamp & Zimmer, 2015; Cheng & Rugg, 2010). These processing characteristics were presumably not initiated by the presentation of the answers to the trivia questions. Taken together, the SMEs in this work turned out to be greater at the beginning of the answer presentation (RewP, P3) and to vanish when it comes to a later time window (LPP).

### Conclusion

In the present work, we specifically targeted the RewP, P3, and LPP because of their relationship with reward processing and found meaningful connections between these ERPs and curiosity. The findings show increased positive amplitudes for all examined ERPs for high compared to low curiosity, suggesting increased reward processing. The SMEs for the RewP and P3 indicate that these reward-related processes contribute to successful memory encoding. According to this first investigation of ERPs during curiosity satisfaction, our findings support the view that curiosity satisfaction resembles reward processing, opening up the investigation of further curiosity-related ERPs studies in future research. Moreover, the present study has shown that not only the state of curiosity but also its satisfaction is an important object of research to better understand the processes underlying curiosity and its similarity to reward. Furthermore, understanding which processes are normally influenced by the satisfaction of curiosity could help to detect clinical conditions that are accompanied by a lack of motivation and dopaminergic circuit activity. In addition, ERPs are valuable tools to explore effects of curiosity due to their high temporal resolution. They allow to track the time course of curiosity-related processing while it unfolds. In summary, our findings suggests that the satisfaction of curiosity with new information is rewarding.

## Data availability statement

The behavioral and ERP data are available at https://osf.io/9k3tv/. The stimuli are available at https://osf.io/he6t9/. The experiment was not preregistered.

## Author contributions

C.R. and M.J.G. designed research, J.C.-D. and M.J.G. performed research, T.R. analysed the data, K.C.J.E. and J.C.-D. assisted and contributed to data analysis, A.M. and M.J.G. supervised the project. T.R. drafted the paper, T.R., A.M., K.C.J.E. and M.J.G. interpreted and wrote the paper, and all other authors gave comments on the paper.

## Acknowledgements and funding information

This work was supported by a Wellcome Trust and Royal Society Sir Henry Dale Fellowship (211201/Z/18/Z) awarded to M.J.G. and a German Research Foundation Research Fellowship (442588275) awarded to K.C.J.E. For the purpose of Open Access, the authors have applied a CC-BY public copyright licence to any Author Accepted Manuscript version arising from this submission. The authors thank Gerrit Höltje (Experimental Neuropsychology Department, Saarland University) for helping during the statistical analysis and writing.

